# Elevated SARS-CoV-2 Antibodies Distinguish Severe Disease in Early COVID-19 Infection

**DOI:** 10.1101/2020.12.04.410589

**Authors:** Natalie S. Haddad, Doan C. Nguyen, Merin E. Kuruvilla, Andrea Morrison-Porter, Fabliha Anam, Kevin S. Cashman, Richard P. Ramonell, Shuya Kyu, Ankur Singh Saini, Monica Cabrera-Mora, Andrew Derrico, David Alter, John D. Roback, Michael Horwath, James B. O’Keefe, Henry M. Wu, An-Kwok Ian Wong, Alexandra W. Dretler, Ria Gripaldo, Andrea N. Lane, Hao Wu, Saeyun Lee, Mindy Hernandez, Vanessa Engineer, John Varghese, Sang Le, Iñaki Sanz, John L. Daiss, F. Eun-Hyung Lee

**Affiliations:** Division of Pulmonary, Allergy, Critical Care & Sleep Medicine, Department of Medicine, Emory University, Atlanta, GA, 30322, USA; MicroB-plex Inc, Atlanta, GA, 30332.; Division of Rheumatology, Department of Medicine, Emory University, Atlanta, GA, 30322, USA; Lowance Center for Human Immunology, Emory University, Atlanta, GA, 30322, USA; Department of Pathology and Laboratory Medicine, Emory University, Atlanta, GA, 30322, USA; Division of Primary Care, Department of Medicine, Emory University, Atlanta, GA, 30322, USA; Division of Infectious Diseases, Department of Medicine, Emory University, Atlanta, GA, 30322, USA; Infectious Disease Specialists of Atlanta, Decatur, GA, 30033, USA; Department of Biostatistics and Bioinformatics, Emory University Atlanta, GA 30322, USA

## Abstract

**Background:** SARS-CoV-2 has caused over 36,000,000 cases and 1,000,000 deaths globally. Comprehensive assessment of the multifaceted anti-viral antibody response is critical for diagnosis, differentiation of severe disease, and characterization of long-term immunity. Initial observations suggest that severe disease is associated with higher antibody levels and greater B cell/plasmablast responses. A multi-antigen immunoassay to define the complex serological landscape and clinical associations is essential.

**Methods:** We developed a multiplex immunoassay and evaluated serum/plasma from adults with RT-PCR-confirmed SARS-CoV-2 infections during acute illness (N=52) and convalescence (N=69); and pre-pandemic (N=106) and post-pandemic (N=137) healthy adults. We measured IgA, IgG, and/or IgM against SARS-CoV-2 Nucleocapsid (N), Spike domain 1 (S1), receptor binding domain (S1-RBD) and S1-N-terminal domain (S1-NTD).

**Results:** To diagnose infection, the combined [IgA+IgG+IgM] or IgG for N, S1, and S1-RBD yielded AUC values −0.90 by ROC curves. From days 6-30 post-symptom onset, the levels of antigen-specific IgG, IgA or [IgA+IgG+IgM] were higher in patients with severe/critical compared to mild/moderate infections. Consistent with excessive concentrations of antibodies, a strong prozone effect was observed in sera from severe/critical patients. Notably, mild/moderate patients displayed a slower rise and lower peak in anti-N and anti-S1 IgG levels compared to severe/critical patients, but anti-RBD IgG and neutralization responses reached similar levels at 2-4 months.

**Conclusion:** This SARS-CoV-2 multiplex immunoassay measures the magnitude, complexity and kinetics of the antibody response against multiple viral antigens. The IgG and combined-isotype SARS-CoV-2 multiplex assay is highly diagnostic of acute and convalescent disease and may prognosticate severity early in illness.

**One Sentence Summary:** In contrast to patients with moderate infections, those with severe COVID-19 develop prominent, early antibody responses to S1 and N proteins.

## Introduction

Since SARS-CoV2 was identified in Wuhan, China in December 2019, the outbreak has evolved into a global pandemic with 36,500,000 cases and over 1,000,000 fatalities as of October 2020. Nasopharyngeal RT-PCR is the diagnostic gold standard but varies widely(*1, 2*) due to timing and quality of sample acquisition. Serologic assays can fill gaps especially for asymptomatic infections and for diagnosis later in disease course even when RNA is undetectable(*3*). Although most patients do well, a small subset succumbs to severe life-threatening disease. Anti-virals and immunomodulatory therapies can decrease disease duration and improve mortality(*4, 5*). Biomarkers to identify at-risk patients may provide opportunities to intervene earlier and improve clinical outcomes.

### SARS-CoV-2 Serology

Characterization of anti-SARS-CoV-2 antibody production is essential for identifying COVID-19 immunity(*6–9*). However, the dynamics of the antibody response and its dependence on clinical severity are additional considerations for interpreting serological results. For example, the evolution of B cell responses to SARS-CoV-2 does not always follow the model of IgM preceding IgG. Instead, many SARS-CoV-2-infected patients demonstrated a simultaneous rise in IgM and class-switched IgG(*10–12*). Antibodies to SARS-CoV-2 spike receptor binding domain (S1-RBD) showed correlation with viral neutralization(*13*) but other immunogenic viral proteins include: S1 and S2 subunits of the spike protein, spike N-terminal domain (S1-NTD), and the nucleocapsid protein (N)(*14*). The spike protein mediates viral entry into the host cell through the S1 subunit that binds ACE2 receptors on host cells and S2 that is involved in fusion. Monoclonal neutralizing antibodies have been generated against the S trimer, RBD, and NTD domains(*15–18*). However, antibody-based diagnostics assays can utilize antigens with neutralizing and non-neutralizing epitopes; thus, we examined anti-S1, anti-S1-RBD, anti-S1-NTD and anti-N antibodies.

### Higher antibody levels in severe/critical disease

The host response to SARS-CoV-2 varies dramatically among individuals. Anti-SARS-CoV-2 antibodies are detectable in patients who have recovered from infection; the magnitude and kinetics of serological responses correlate with clinical severity(*3, 19–23*). Reports on SARS-CoV-1 infections showed a paradoxical rise of neutralizing antibodies in critically ill patients(*24*). Similarly, in SARS-CoV-2, the induction of virus-specific antibody responses in severe disease trend higher than in non-severe patients 8-20 days post-symptom onset (DPSO)(*10, 22*). However, there was significant overlap of antibody levels between the two groups such that it could not distinguish mild from severe illness. Other studies corroborated these findings with binding and neutralizing antibodies, but the numbers were small and timing of sample collection was late in disease course(*25–27*).

In this study, we developed a sensitive anti-SARS-CoV-2 immunoassay using pre-pandemic controls and RT-PCR-confirmed, SARS-CoV-2-infected patients and observed higher antibody levels earlier in patients with severe/critical infections compared to those with mild/moderate disease. This unexpected difference in the speed and magnitude of humoral responses may reflect important differences in the immune response and provide early signs for patients who might benefit from targeted immunomodulatory interventions.

## Results

### Acute and convalescent SARS-CoV-2-infected patients

353 adults were enrolled for this study: 110 RT-PCR-confirmed, SARS-CoV-2-infected patients and 243 healthy adult controls (**Table S1**). The pre-pandemic control group consisted of 106 adults whose samples were collected between July 2016 and February 2019. The post-pandemic control group, who had no known exposure to SARS-CoV-2, consisted of 137 adults whose blood samples were collected between March and June 2020.

We enrolled 69 SARS-CoV-2-infected patients after recovery from illness and at least 31 DPSO. This convalescent population provided serum samples used for development and validation of the immunoassay (**Figs.1,3**). Nineteen patients had severe/critical disease (10 severe; 9 critical) and 50 suffered mild/moderate infections (34 mild; 16 moderate) as defined by the NIH criteria(*28*).

**Figure 1.**
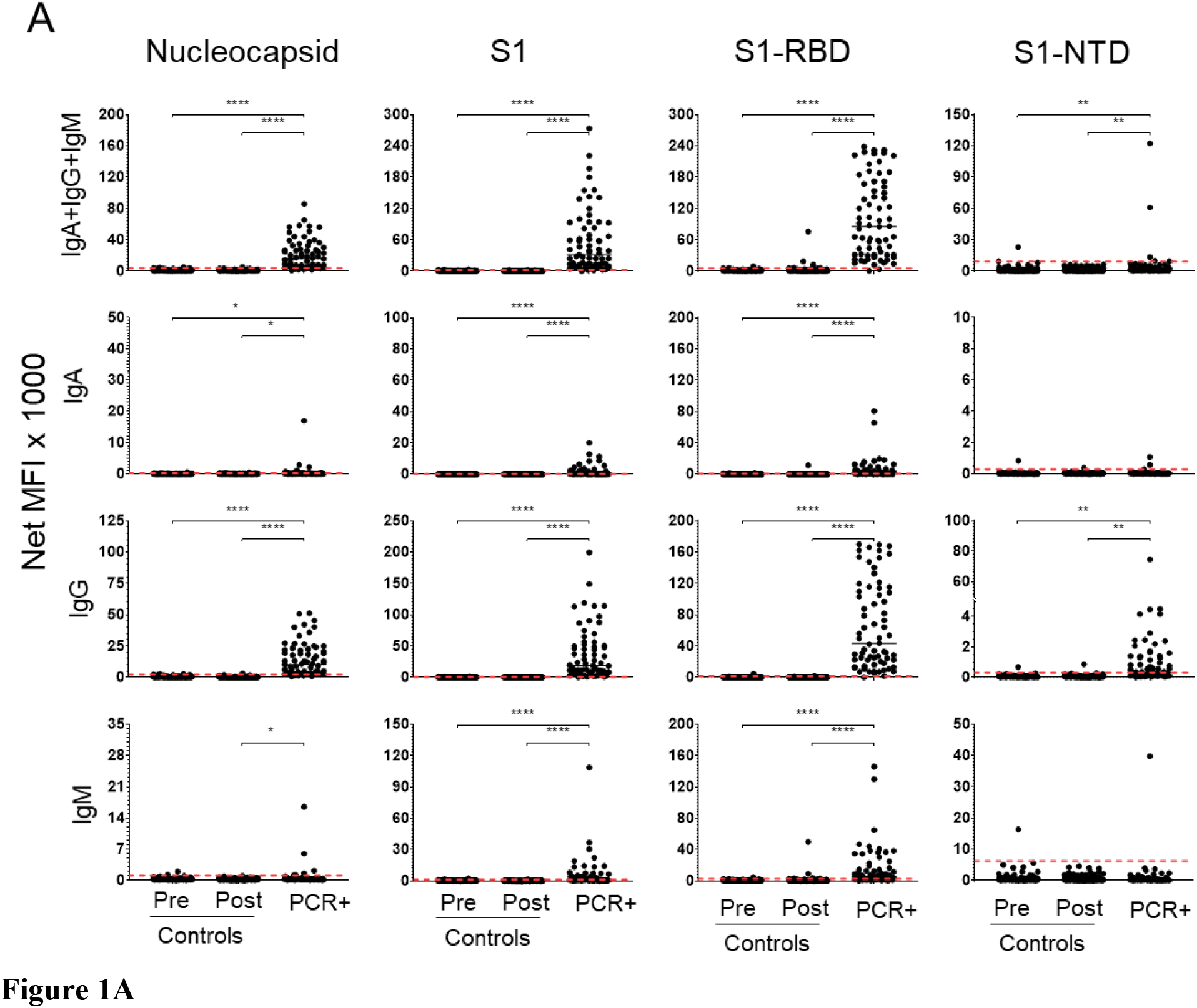
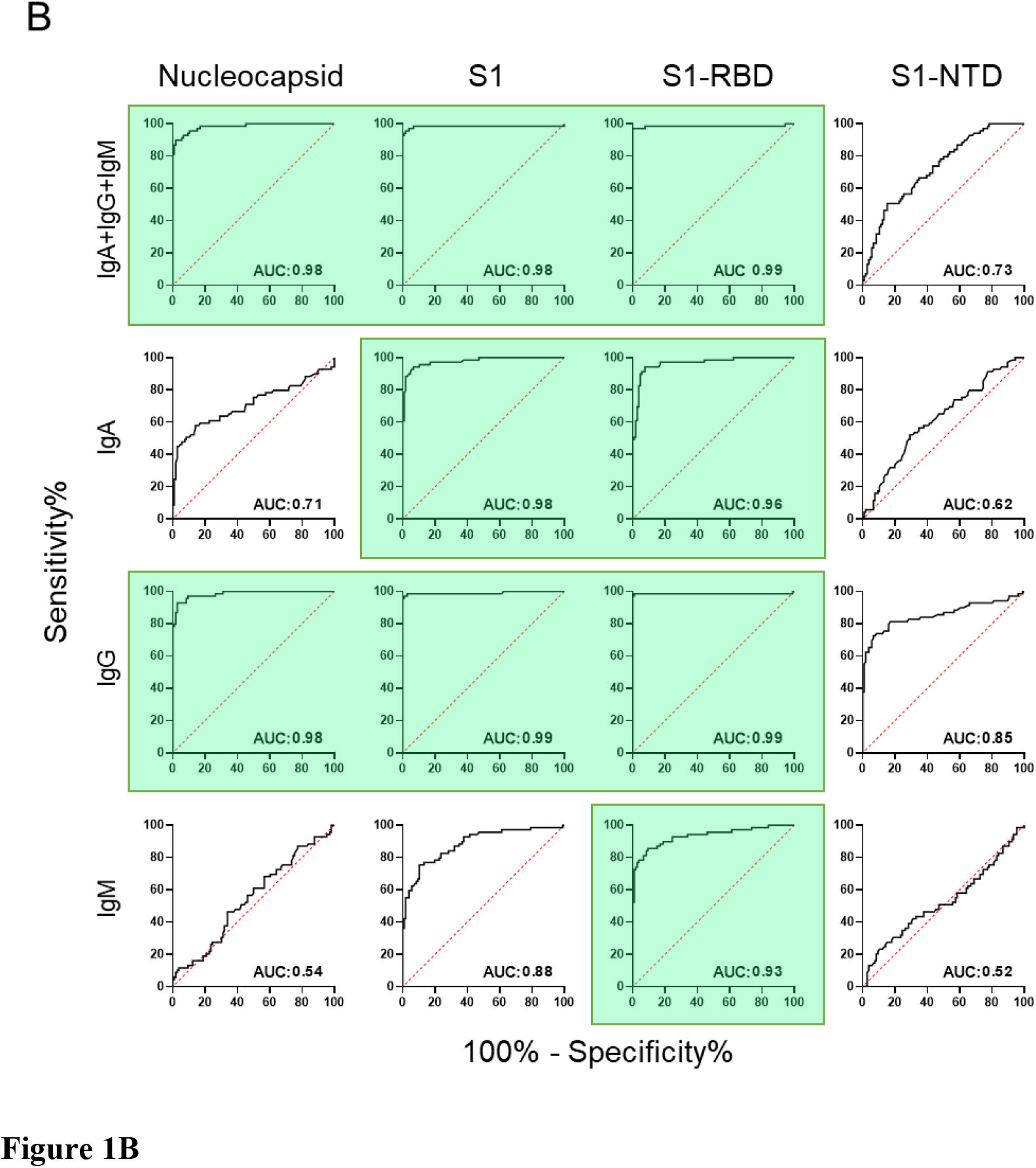
Serum antibody responses distinguish convalescent, RT-PCR-confirmed, SARS-CoV-2-infected patients from controls. **A.** Sera were collected from: i) 106 healthy subjects prior to March 2019 (Pre-pandemic Controls); ii) 137 healthy subjects between March and June 2020 (Post-pandemic Controls) and iii) 69 RT-PCR-confirmed convalescent patients (PCR+) at least 31 days post-symptom onset (DPSO). Each serum sample was diluted 1:500 and then measured for the presence of anti-SARS-CoV-2 antibodies: N, S1, S1-RBD and S1-NTD. Secondary antibodies were a combination of PE-conjugated anti-IgA, anti-IgG, and anti-IgM (top row) or each individually (rows 2-4, respectively). Net MFI is the mean fluorescence intensity with background subtracted. Horizontal, red, dashed lines indicate the diagnostic cut-off (C0) values determined as the mean of the pre-pandemic population net MFI + 3 standard deviations. Significant differences between populations were determined by one-way ANOVA and Tukey’s tests: * p<0.05; ** p<0.01; *** p<0.001; **** p<0.0001; p>0.05 not shown. **B)** Receiver Operating Characteristic (ROC) curves were used to compare diagnostic potential of each SARS-CoV-2 antigen and antibody combination. The 69 RT-PCR-confirmed convalescent patients were compared against the 106 pre-pandemic healthy control subjects. Diagonal, dashed, red lines represent the predicted behavior when the test has no diagnostic value. The area-under-the-curve (AUC) measurement provides an index of diagnostic potential. AUC near 0.5 suggest no diagnostic value and AUC near 1.0 indicate strong diagnostic potential. Antigen:isotype combinations that yielded AUC values greater than 0.90 are highlighted in green.

We enrolled 52 SARS-CoV-2-infected patients during their acute infections (<31 DPSO). Eleven patients were in both the acute and convalescent populations. There were 39 patients classified as severe/critical (3 severe, 36 critical) and 13 as mild/moderate (13 mild, 0 moderate).

### SARS-CoV-2 multiplex immunoassay

Using sera from 69 convalescent patients, we developed a SARS-CoV-2 multiplex immunoassay to detect the combined [IgA+IgG+IgM] or individual IgA, IgG, or IgM responses specific for four viral antigens nucleocapsid (N), S1, S1-RBD, and S1-NTD (**Fig.1A**). Cut-off values (C0) were set at the mean plus three standard deviations for each antigen:secondary antibody combination using the pre-pandemic control population (N=106; **Fig.1A**). Receiver operating characteristic (ROC) curves, where the area-under-the-curve (AUC) provides an estimate of diagnostic potential, are presented in **Fig.1B**.

For the combined isotype [IgA+IgG+IgM] detection, responses to three of the candidate antigens, N, S1 and S1-RBD, had significant diagnostic potential (AUC≥0.98); the response to S1-NTD was relatively poor (AUC=0.73) (**Fig.1B**). Overall, SARS-CoV-2-specific IgM antibodies were poor predictors of convalescent infection; AUC values were low for IgM S1-NTD (AUC=0.52) and N (AUC=0.54), although, IgM levels for S1 (AUC=0.88) and S1-RBD (AUC=0.93) were potentially useful. IgA antibodies were fairly strong predictors of convalescent infection with AUC values greater than 0.95 for S1 and S1-RBD. However, IgG responses had the greatest diagnostic potential for each of the three antigens N, S1 and S1-RBD (AUC≥0.98); anti-S1-NTD IgG was lower (AUC=0.85). In the convalescent population, IgG responses against the S1 or the S1-RBD antigens were specific and sensitive for identifying patients who had experienced SARS-CoV-2 infections.

### Prozone effect in patients with severe disease

To assess sample limitations in the assay, we titrated ten serum samples: four from patients with severe/critical disease, three from patients with mild/moderate disease, and three from pre-pandemic controls. We observed the expected log-linear dose-response curves in the mild/moderate samples, but we detected bell-shaped binding curves in the severe/critical groups, indicative of the prozone effect in the antigen:antibody interactions for N, S1, and S1-RBD (**Fig.2A**). Subsequent analyses were performed with samples diluted 1:500 to maximize the measurement of low-responding mild/moderate patients while minimizing the potential impact of the prozone effect in severe/critical patients.

**Figure 2.**
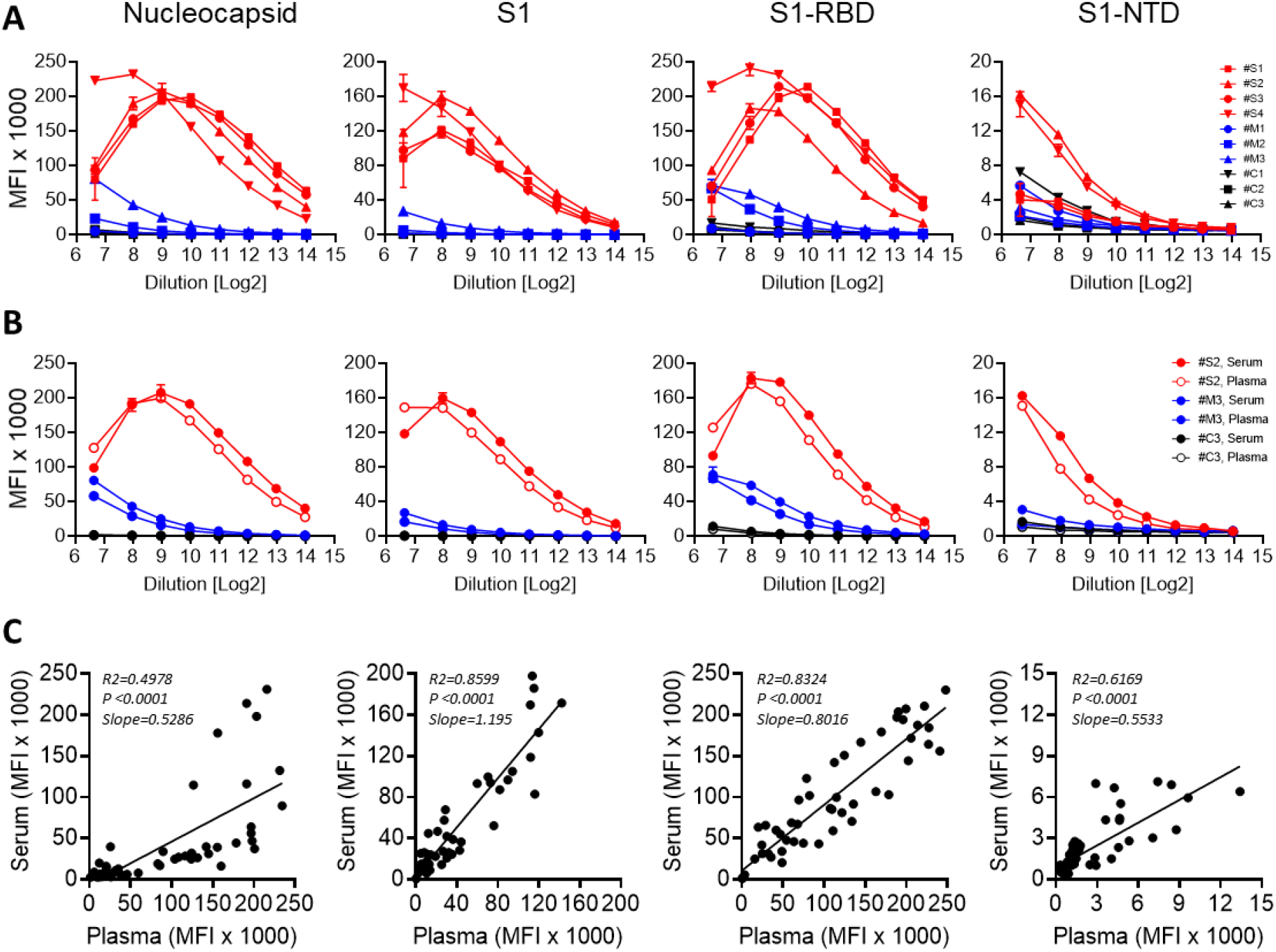
Prozone effect in sera from severe/critical infections and comparison of the assay using serum and plasma. **A)** Titration of the assay using sera from four severe (red), three mild (blue), and three healthy pre-pandemic controls (black) for [anti-IgG + anti-IgA + anti-IgM] specific for the N, S1, S1-RBD, and S1-NTD antigens. Sample dilutions range from 1:100 (2^6.64^) to 1:16,384 (2^14^). **B)** Comparison of plasma (open circles) and serum (closed circles) in the multiplex assay for N, S1, S1-RBD, and S1-NTD in one representative severe, mild, and healthy control. **C)** Correlation of antibody titers in 50 matched serum and plasma samples measured at a 1:500 dilution.

### Antibody concentrations in serum of SARS-CoV-2-infected patients

To estimate a quantity of antibodies using monoclonal antibody standards, we measured anti-S1 IgG concentrations were as high as 45 μg/mL in sera from mild/moderate patients and 100-400 μg/mL in sera from severe/critical patients. N-specific antibodies ranged from 1-20 μg/mL and 17-49 μg/mL in the same groups.

### Serum and plasma samples yield similar results

Nearly identical binding curves from a small set of paired serum and plasma samples were observed including the prozone effect (**Fig.2B**). Paired serum and plasma samples from 50 subjects yielded similar MFI values with particularly high correlations (R^2^>0.83) for S1 and S1-RBD antigens (**Fig.2C**). Measurement of antibody levels at a 1:500 dilution resulted in comparable values for serum and plasma samples from severe/critical and mild/moderate patients.

### Elevated levels of anti-SARS-CoV-2 antibodies in sera from convalescent patients with severe/critical infections compared to those with mild/moderate disease

We observed a correlation between disease severity and the magnitude of the antibody response when the convalescent COVID-19 patients were divided into mild/moderate (N=50) and severe/critical (N=19) cohorts. (**Fig.3**). The combined isotype response and IgG alone against S1 and S1-RBD resolved the two populations significantly (AUC>0.9; Fig.S1). The IgA and IgM responses to the viral antigens were less discriminating of the two groups (AUC<0.86). N and S1-NTD were not useful antigens for separating the severe/critical from the mild/moderate groups (AUC<0.86), except for S1-NTD IgG (AUC>0.91). Elevated antibody levels in convalescent severe/critical patients suggested that this striking difference may be evident earlier during infection.

**Figure 3.**
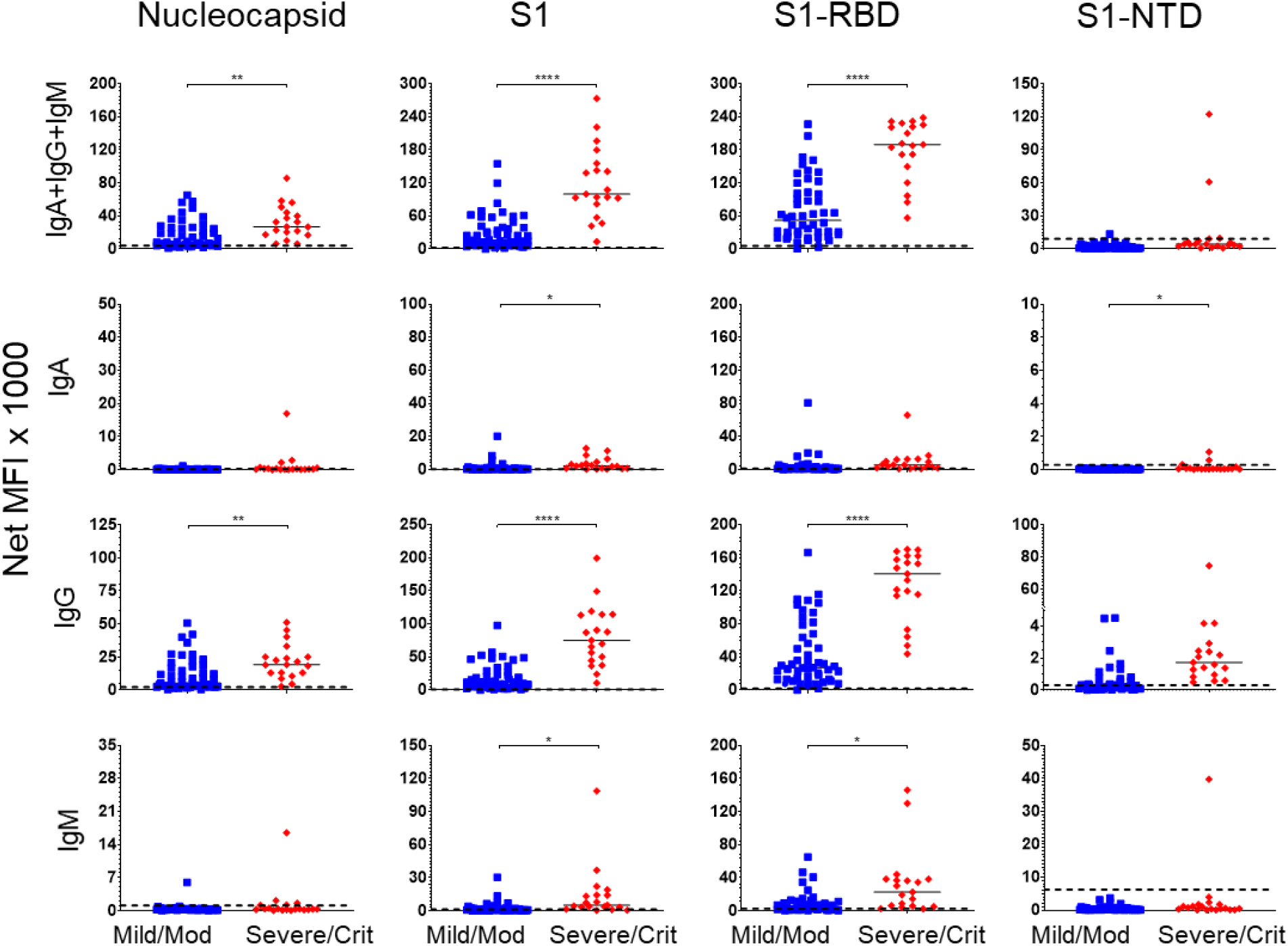
Convalescent patients who had suffered severe/critical infections have higher anti-SARS-CoV-2 antibody levels than patients who had suffered mild/moderate infections. The serum responses of the 69 convalescent SARS-CoV-2-infected patients in Fig. 1 were assorted according to the severity of their infections: mild/moderate (N=50, blue squares) and severe/critical (N=19, red diamonds). Median values for each group are presented as black, horizonal bars. Horizontal, black, dashed lines indicate the diagnostic cut-off (C0) values determined by the Pre-pandemic controls. Significance determined by a two-tailed t-test: * p<0.05; ** p<0.01; *** p<0.001; **** p<0.0001; p>0.05 not shown.

### Elevated serum antibody levels early in infection of severe/critical patients

Among the 110 infected patients, sera were collected from 50 patients within 6-30 DPSO (39 severe/critical, 11 mild/moderate infections; median draw: 13 DPSO). As in the convalescent population, S1 and S1-RBD antigens yielded the greatest diagnostic potential (AUC≥0.94) in combined or individual isotypes (**Fig.S2**). In contrast to the convalescent serum samples, measurement of IgA, IgG or the combined isotypes against N exhibited potential utility (AUC≥0.90). Furthermore, in acute illness, IgA specific to N, S1 and S1-RBD as well as IgM to S1 and S1-RBD yielded significantly better AUC values than they did in the convalescent patient population (**Fig.1B**). Thus, IgA, IgM or IgG could potentially be used for detection of early infection.

Separating the SARS-CoV-2-infected patients by disease severity reveals striking differences between the mild/moderate and severe/critical populations (**Fig.4**). As in the convalescent population, the strongest predictors of severe/critical infections were the combined isotypes and IgG responses to the S1 and S1-RBD (AUC>0.91, **Fig.S3**). However, in the interval 6-30 DPSO, anti-S1, anti-S1-RBD, and anti-N IgG or the combined isotypes showed comparably high AUC values (**Fig.4**, **Fig.S3**). IgG and IgA, but not IgM, predominated in the patients with early severe/critical disease suggesting that early class-switched responses are evident in severe/critical infections.

**Figure 4.**
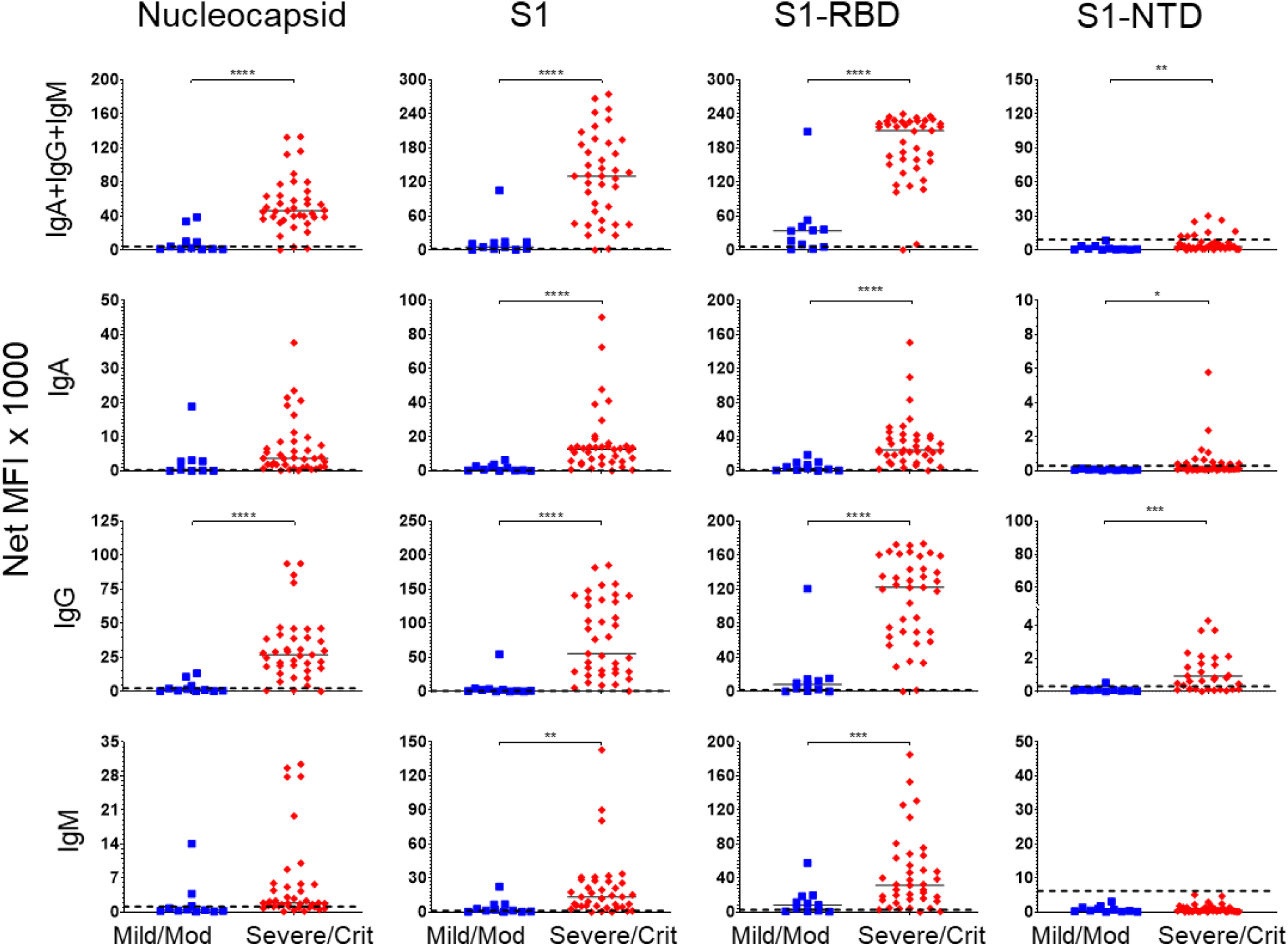
Higher SARS-CoV-2 antibody levels in severe/critical compared to mild/moderate infections during acute illness (≤30 DPSO). Serum antibody levels were compared in 39 severe/critical patients (red diamonds) and 11 mild/moderate patients (blue squares) using samples drawn 6 to 30 DPSO. Horizontal, red, dashed lines indicate the C0 values (from Fig. 1). Horizontal, black, dashed lines indicate the diagnostic cut-off (C0) values determined by the pre-pandemic population. Significance determined by a two-tailed t-test: * p<0.05; ** p<0.01; *** p<0.001; **** p<0.0001; p>0.05 not shown.

### Kinetics of anti-SARS-CoV-2 antibodies in severe/critical and mild/moderate groups

To determine the kinetics, magnitude, and durability of serum antibody responses, we analyzed sera from 165 samples that included 53 severe/critical and 57 mild/moderate patients ranging from 2-150 DPSO (**Fig.5**). There was an early significant increase in antibody responses in the severe/critical groups for nearly all isotypes specific for N, S1, S1-RBD and S1-NTD. The only exception was IgM for S1-NTD (Table S2). At later time points >90 days, the antibody responses were similar between the two groups. These results suggest that for the severe/critical cohort, anti-SARS-CoV-2 antibodies rise rapidly while in the mild/moderate group antibodies rises slower. In summary, disease severity can affect magnitude, isotype, specificity, and potentially durability of antibody responses.

**Figure 5.**
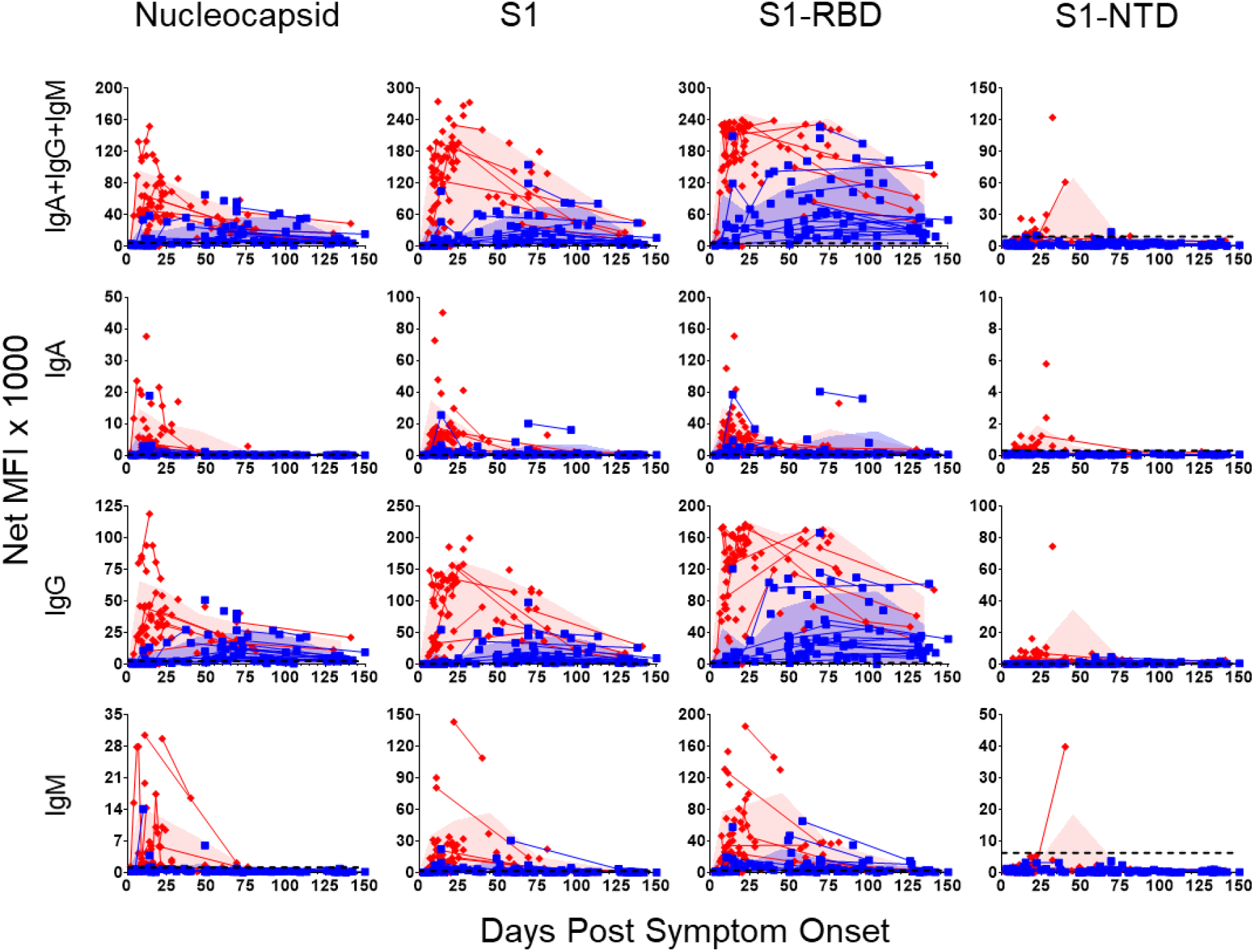
Comparison of longitudinal antibody responses in severe/critical and mild/moderate SARS-CoV-2 patients over 150 days. Serum samples were collected from patients with mild/moderate (86 samples from 57 patients, blue squares) or severe/critical (79 samples from 53 patients, red diamonds) SARS-CoV-2 infections from days 2-150 DPSO. Lines connect multiple samples drawn from the same patient. Shaded blue and red areas represent the mean plus standard deviation of mild/moderate or severe/critical patient groups, respectively. during 0-15, 16-30, 31-60, 61-90, 91-120,121-150 DPSO and plotted at the midpoint of each time interval. Number of samples used at each time interval: mild/moderate N = 13, 6, 21, 21, 11, 14; severe/critical N = 33, 23, 9, 9, 0, 4. Data point was skipped for the severe/critical group at the 91-120 DPSO interval since only one sample was drawn during that timeframe.

### Viral neutralization detected in serum with high SARS-CoV-2 antibody levels

Neutralizing antibodies are capable of blocking the interaction between the SARS-CoV-2 RBD and the ACE2 receptor on the surface of target cells. To determine whether sera from infected patients had SARS-CoV-2 neutralizing activity, we tested a representative subset of 19 plasma samples drawn within 30 DPSO from 12 severe/critical patients, 4 mild/moderate patients, and 3 healthy pre-pandemic subjects using a surrogate viral neutralization test (sVNT) (*29*). Sera from patients in the severe/critical group demonstrated inhibition of ACE2 receptor-binding activity by HRP-conjugated S1-RBD, achieving 50% reductions at several hundred-fold dilutions (Log_10_ sIC50 median = 2.48), compared to little or no measurable activity (Log_10_ sIC50 <0.5) in the mild/moderate and healthy control groups (**Fig. 6A**). Using CR3022 (InvivoGen) and 6G9 (Amsbio) as controls, we also titrated the S1-RBD antibodies and found concentrations of 0-45 ug/mL and 100-400 ug/mL in mild/moderate and severe/critical cohorts, respectively. N-specific antibodies ranged from 1-20 ug/mL and 17-49 ug/mL in mild/moderate and severe/critical groups, respectively. In a second population of severe/critical (N=15) and mild/moderate (N=10) patients in **Fig. 6B** (p < 0.0001), we validated assessed neutralization using the sVNT. In plasma/serum samples collected in the convalescent stage of the infection (days 37 to 140), the neutralizing titers in severe/critical patients (N=8) decline slightly while those measured in the mild/moderate patients (N=23) rise substantially to near equivalence with the severe/critical population.

**Figure 6.**
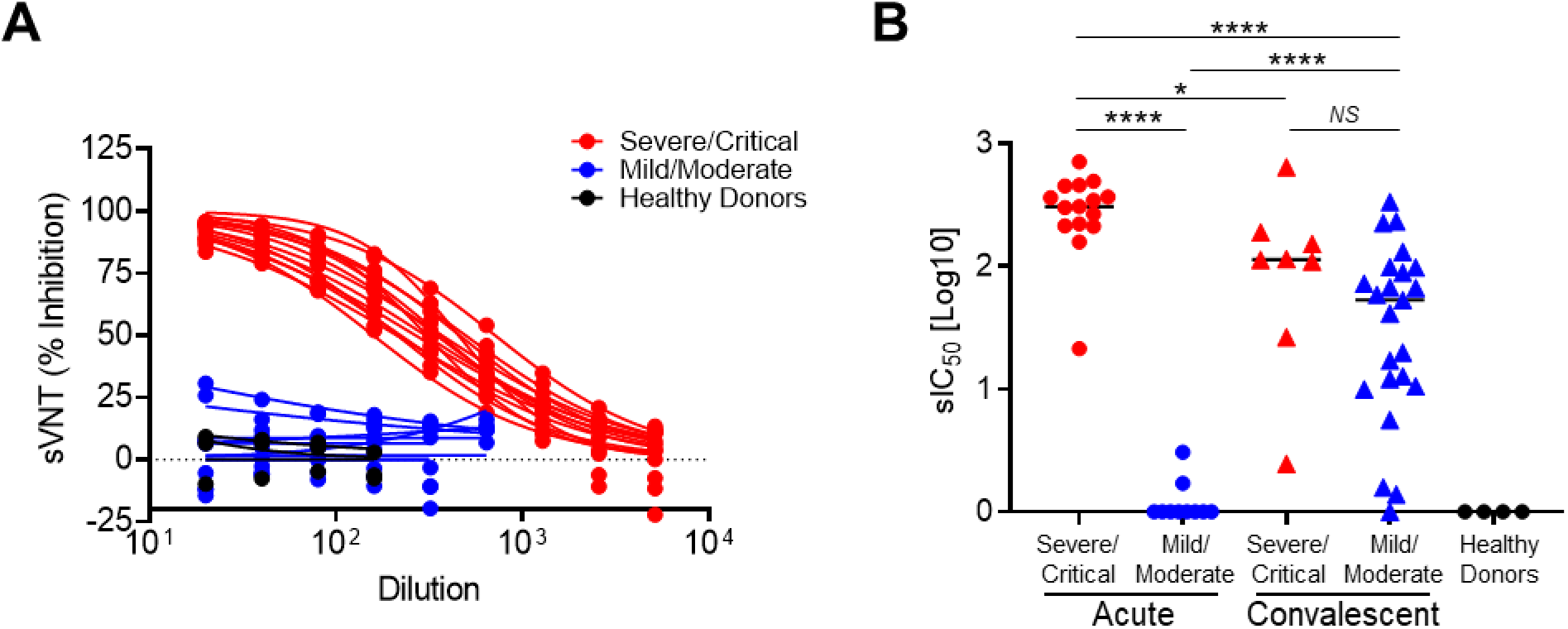
Quality of SARS-CoV-2 antibodies in severe/critical and mild/moderate during acute infection and convalescence. **A)** Dose-response curves of sera from patients with severe/critical (red lines), or mild/moderate (blue lines) SARS-CoV-2 infections and healthy controls (black lines) using the GenScript surrogate Virus Neutralization Test (sVNT) that measures blockade of the interaction between the immobilized ACE2 receptor and soluble HRP-conjugated S1-RBD by sample-borne antibodies. sVNT (% Inhibition) is the percentage reduction of the ACE2 receptor:HRP-S1-RBD interaction in the absence of an inhibitor. **B)** Serum dilution yielding 50% reduction of the uninhibited response in the sVNT sIC_50_[Log10] observed in severe patients (red dots, triangles); mild patients (blue dots, triangles) and healthy controls (black dots) during acute illness and convalescence. The significance determined by a two-sided t-test: not significant (NS) p>0.05; * p<0.05; ** p<0.01; *** p<0.001; **** p<0.0001.

## Discussion

In order to visualize the comprehensive landscape of SARS-CoV-2 humoral immunity, we developed a sensitive, multiplex immunoassay for detection of acute infection and previous viral exposure. This immunoassay detects individual or combined IgA, IgG, and IgM antibodies against viral antigens N, S1, S1-RBD, and S1-NTD. Due to its wide dynamic range, the assay can also distinguish severe/critical from mild/moderate infections and potentially prognosticate severity early in infection. However, we observed in severe/critical sera an unexpected prozone effect that may have been underappreciated in prior studies. Additionally, neutralizing antibodies correlated with viral-specific antibody titers, particularly in severe/critical patients early in disease and rose in mild/moderate patients only after recovery. For diagnosis of acute and convalescent infections, the combined-isotype or IgG immunoassays for anti-S1-RBD performed best, although anti-N and anti-S1 provided confirmation. To distinguish mild/moderate from severe/critical infections, measurement of combined isotypes or IgG specific for S1-RBD, S1, or N had high predictive values (AUC≥0.90). In particular, anti-S1-RBD was excellent for both diagnosis and prognosis of severity.

### Differences in longitudinal antibody responses and disease severity

Magnitude and kinetics of SARS-CoV-2 antibody responses differed with disease severity. Severe/critical patients (nine of whom died) reached very high titers against N, S1, and S1-RBD at 6-20 DPSO. These early responses consisted of not only IgM but primarily of class-switched IgG and IgA antibodies.

The decline in antibody levels occurred at different rates for several antigen:isotype combinations. The levels of IgG, IgA, and IgM against N and S1 decline rapidly after recovery in the severe/critical group; thus, they would not useful for convalescent diagnosis. In contrast, the mild/moderate patients had slower rises of antibody titers, mostly after 30 DPSO, and slower declines evident only after 3-4 months. Patients with severe/critical disease have increased ASC expansions early in infection(*12, 30, 31*) associated with higher antibody levels compared to those observed in mild/moderate patients. Together, these results suggest different sources of B cells and ASC in the severe/critical and mild/moderate infection groups.

### Prozone effect and previous underestimation of SARS-CoV-2 antibodies

We could discriminate serum titers between severe/critical and mild/moderate patients early in illness due to the sensitivity of this platform. It is well known that the prozone (or hook) effect occurs when an excess of antigen or antibody inhibits the formation of cross-linked Ag-Ab complexes(*32–35*). In our assay, this effect was observed at higher serum concentrations demonstrating abundant antibody levels in severe/critical patients. Concentrations of anti-S1-RBD IgG rose to 100-400 μg/mL in severe/critical patients by 10 DPSO and approached similar levels in mild/moderates after 30 DPSO. Our results suggest that previous studies may have underestimated titers in patients with severe disease due to the prozone effect. Thus, the need for optimizing performance characteristics for these assays will be essential as we standardize serologic testing.

### Severe/critical and mild/moderate patients display striking differences in their humoral immune responses: clinical significance and immunological basis

Severe/critical patients produce a vigorous early antibody response to SARS-CoV-2 comprised primarily of IgG and IgA, and little IgM. In contrast, mild/moderate patients develop a conventional immune response that develops over 4-6 weeks. The striking early response produced by severe/critical patients may enable early detection of the patients most likely to experience poor outcomes.

Conventional models of B cell responses to primary infections include an early extrafollicular response that provides a short-lived IgM wave of antibodies with very low mutation load and antigen affinity. This extrafollicular phase precedes subsequent germinal center reactions that generate memory B cells and plasma cells that secrete higher affinity, isotype-switched antibodies(*36*). It is now evident however that extrafollicular responses also generate isotype-switched antibody responses(*37*), and that this process may indeed be prominent in SARS-CoV-2 infections as recently demonstrated by our groups and others(*12, 38*). Moreover, our work showed that severe infection is strongly associated with intense extrafollicular B cell responses producing IgG and IgA antibodies with very low levels of somatic hypermutation despite the presence of high neutralizing antibody titers(*12*). In keeping with our results, severe COVID-19 infections create profound disruption of germinal center structures(*38*). Combined, the available evidence strongly suggests that early responses to SARS-CoV-2 may be comprised of simultaneous early generation of IgM, IgG and IgA responses with low mutation rates through intense extrafollicular responses and defective germinal center reaction. This profile would be exaggerated in severe cases. Late and post-infection responses would depend on the degree of initial disruption and/or subsequent restoration of proper germinal center responses.

In all, our sensitive, multiplex immunoassay is ideal for diagnosis of acute and convalescent infections and important for predicting severity early in the disease course. We conclude that the virus-specific antibody kinetics vary depending on disease severity and we speculate that specific viral antigens will have implications for long-term humoral protection to SARS-CoV-2 re-infection.

## Materials and Methods

### Institutional Approval for Patient Sample Collection

All blood and tissue samples were collected under the Emory University Institutional Review Board approved protocols. Pre-pandemic samples were collected from healthy adults prior to the pandemic between July 2016 and February 2019. After the start of the pandemic, samples were collected from March-June 2020 from healthy adults with no known exposure to SARS-CoV-2 (**Table S1**). Patients with RT-PCR-confirmed, SARS-CoV-2 infections were enrolled from Emory University hospitals and outpatient facilities. Severity of SARS-CoV-2 infected patients was determined according to NIH guidelines(*28*) (**Table S1**). Patients were categorized based on their overall disease course as mild/moderate or severe/critical. Samples collected in the first 30 DPSO were considered acute and those collected after 30 DPSO were designated convalescent. All samples were processed within 4-12 hours of collection and stored at −80°C for subsequent analysis.

### Selection of Antigens

Four recombinant SARS-CoV-2 antigens were used in this study. The Nucleocapsid protein (N; Cat. No. Z03480; expressed in *E. coli*), the S1 domain (S1; amino acids 16-685; Cat. No. Z03485; expressed in HEK293 cells) of the Spike protein, and the S1-Receptor Binding Domain (S1-RBD; Cat. No Z03483; expressed in HEK293 cells) were purchased from by GenScript. The S1-N-terminal domain (S1-NTD, amino acids 16-318) was custom synthesized by GenScript. Each protein was expressed with an N-terminal His6-Tag to facilitate purification, greater than 85% pure and appeared as a predominant single band on SDS-PAGE analysis.

### Carbodiimide coupling of microspheres to SARS-CoV-2 antigens

*SARS-CoV-2* proteins were coupled to MagPlex® Microspheres of spectrally distinct regions (Luminex; Austin, TX, USA). Coupling was carried out at room temperature following standard carbodiimide coupling procedures. Microspheres were washed once with deionized water and incubated for 20 minutes in the dark in an end-over-end rotator suspension of 0.1 M NaH2PO4 (VWR; Radnor, PA, USA), 5 mg/mL EDC (1-Ethyl-3-(3-dimethylaminopropyl) carbodiimide; Thermo Fisher Scientific; Waltham, MA, USA), and 5 mg/mL Sulfo-NHS (N-hydroxysulfosuccinimide; Thermo Fisher Scientific; Waltham, MA, USA). Activated microspheres were washed twice in 0.05 M MES (2-(N-morpholino) ethanesulfonic acid, Boston Bioproducts, Ashland, MA, USA) and incubated for two hours in the dark, in an end-over-end rotator 500 μL suspension of 0.05 M MES containing 300 nMol/L antigen and 1×10^6^ microspheres (0.15 nMol antigen per 10^6^ microspheres). Coupled microspheres were washed four times in a blocking buffer consisting of 1% BSA (Bovine Serum Albumin; Boston Bioproducts; Ashland, MA, USA), 1X PBS (VWR; Radnor, PA, USA), 0.05 % sodium azide (VWR; Radnor, PA, USA), and 0.02 % Tween-20 (VWR; Radnor, PA, USA). Coupled microspheres were then stored at a concentration of 10^6^ spheres/mL at 4 °C in the same blocking buffer. Concentrations of spheres were confirmed by counting on a Bio-Rad T20 Cell Counter.

### Luminex immunoassays for measurement of anti-SARS-CoV-2 antibodies

Fifty μL coupled microsphere mix was added to each well of clear-bottomed, 96-well, black, polystyrene microplates (Greiner Bio-One North America Inc., Monroe, NC, USA) at a concentration of 1000 microspheres per region per well. All wash steps and dilutions were accomplished using 1% BSA, 1X PBS assay buffer. Except when assay parameters were under development (Fig. 2), serum was assayed at 1:500 dilution and surveyed for antibodies specific for the four candidate antigens. After a one-hour incubation, in the dark, on a plate shaker at 800 rpm, wells were washed five times in 100 μL of assay buffer, using a BioTek 405 TS plate washer. Then, the secondary reagent, PE-conjugated Goat Anti-Human IgA, IgG and/or IgM (Southern Biotech; Birmingham, AL, USA), was added at 3 μg/mL. After 30 minutes of incubation at 800 rpm in the dark, wells were washed three times in 100 μL of assay buffer, resuspended in 100 μL of assay buffer, and analyzed using a Luminex FLEXMAP 3D® instrument (Luminex; Austin, TX, USA) running xPonent 4.3 software. Median Fluorescent Intensity (MFI) using combined or individual detection antibodies (anti-IgA/anti-IgG/anti-IgM) was measured using the Luminex xPONENT software. The background value of assay buffer was subtracted from each serum sample result to obtain Median Fluorescent Intensity minus Background (MFI-B or Net MFI).

### Determination of Cut-off (C_0_) Values

Determination of cut-off values was based on quantitative evaluation of the pre-pandemic healthy controls. The C_0_ for each antigen/secondary combination was defined as the mean plus three standard deviations of the 106 pre-pandemic samples.

### Surrogate Virus Neutralization Test (sVNT) and antibody concentrations

Circulating antibodies that block the interaction between ACE2 receptor and SARS-CoV-2 RBD were measured using a recently described surrogate Virus Neutralization Test (sVNT)(*29*), following manufacturer’s recommendations. Briefly, plasma or serum samples were diluted with sample dilution buffer, followed by incubation with HRP-conjugated SARS-CoV-2 RBD (HRP-S1-RBD); the mixtures were then transferred to 96-well plates coated with immobilized, recombinant ACE2 receptor. Following washing and TMB development, the optical density values at 450 nm were measured. The level of inhibition, i.e., the reduction of binding of HRP-RBD to the immobilized ACE2 receptor, was calculated as: % Inhibition = (1-(OD sample/OD negative controls)) x 100. Surrogate IC50 (sIC50) values are the dilutions of sample that yielded 50% reductions in the sVNT assay. Monoclonal antibodies CR3022 (human IgG1 anti-S1-RBD(*39*); InvivoGen) and 6G9 (human:mouse chimeric IgG anti-N; Amsbio) were used as calibrators for SARS-CoV-2 antibody concentrations.

### Data analysis

ANOVA, Receiver Operating Characteristic (ROC) curves were generated and Area-Under-the-Curve (AUC) values were calculated. Comparisons among and between groups were made using one-sided ANOVA or two-sided t-tests using GraphPad Prism.

## Supported by

NIH: NIAID: 3P01AI125180-05S1, 1R01AI121252, P01A1078907, U01AI045969, U19AI109962, U54CA260563, T32-HL116271-07, NIGMS 2T32GM095442;

The Marcus Foundation; CDC: Contract #200-215-88233.

## Supplemental Figures and Tables

**Figure S1.**
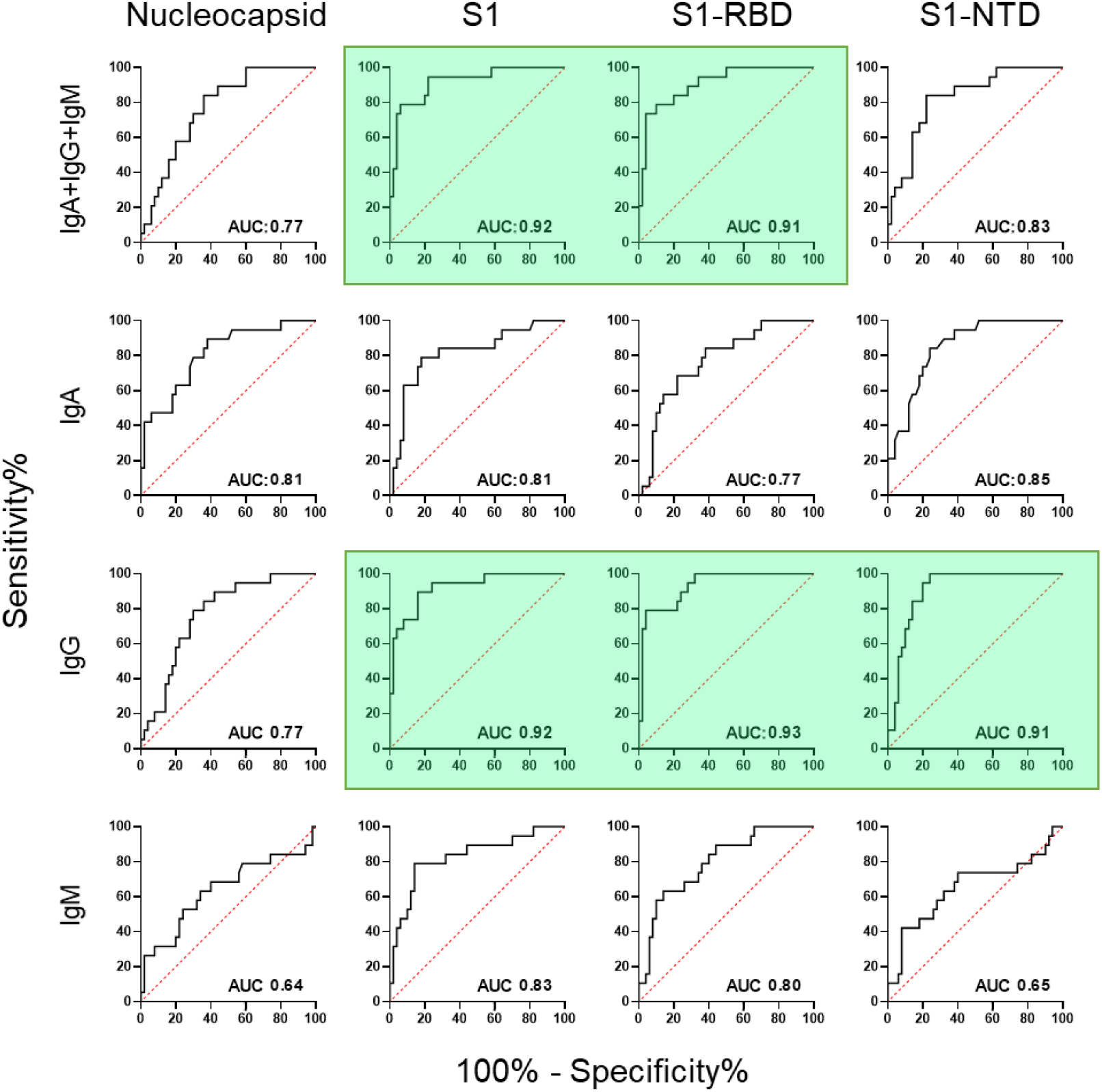
ROC curves distinguish severe/critical and mild/moderate SARS-CoV-2 infections during convalescence. ROC curves were used to compare serum SARS-CoV-2 antibody responses between the 19 severe/critical and 50 mild/moderate infected patients (from Fig. 3) during convalescence. Diagonal, dashed, red lines represent the predicted behavior when the test has no diagnostic value. The area-under-the-curve (AUC) measurements provides an index of potential diagnostic utility: AUC near 0.5 suggest no diagnostic value; AUC values near 1.0 indicate strong diagnostic potential. Antigen:isotype combinations that yielded AUC values greater than 0.90 are highlighted in green.

**Figure S2.**
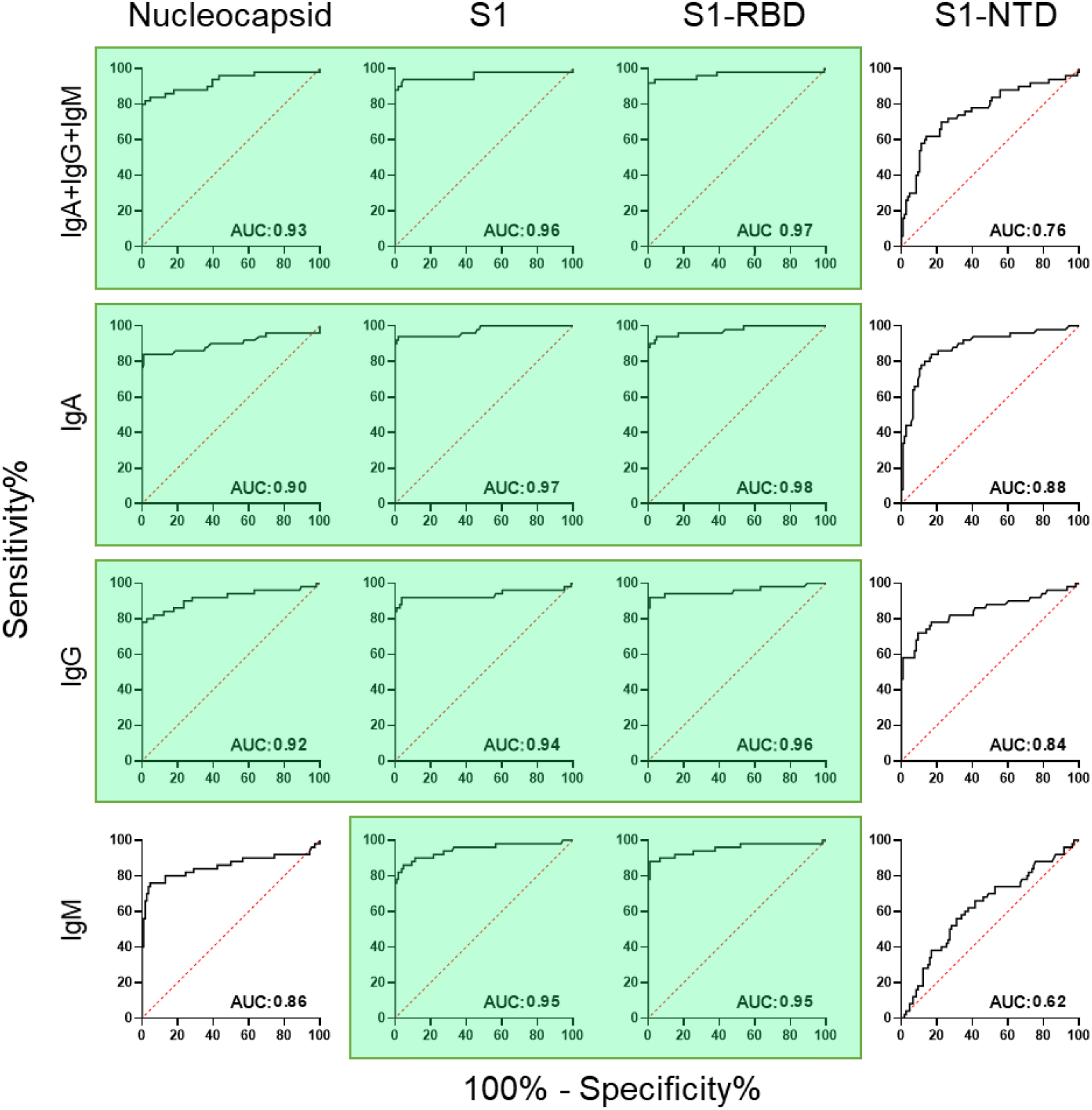
ROC curves assess diagnostic potential of SARS-CoV-2 antibody levels during acute illness. ROC curves compare 50 acutely infected SARS-CoV-2 patients (6-30 DPSO) against 106 pre-pandemic controls. Diagonal, dashed, red lines represent the predicted behavior when the test has no diagnostic value. The area-under-the-curve (AUC) measurements provide an index of diagnostic utility where AUC near 0.5 suggest no diagnostic value and AUC near 1.0 indicate strong diagnostic potential. Antigen:isotype combinations that yielded AUC values greater than 0.90 are highlighted in green.

**Figure S3.**
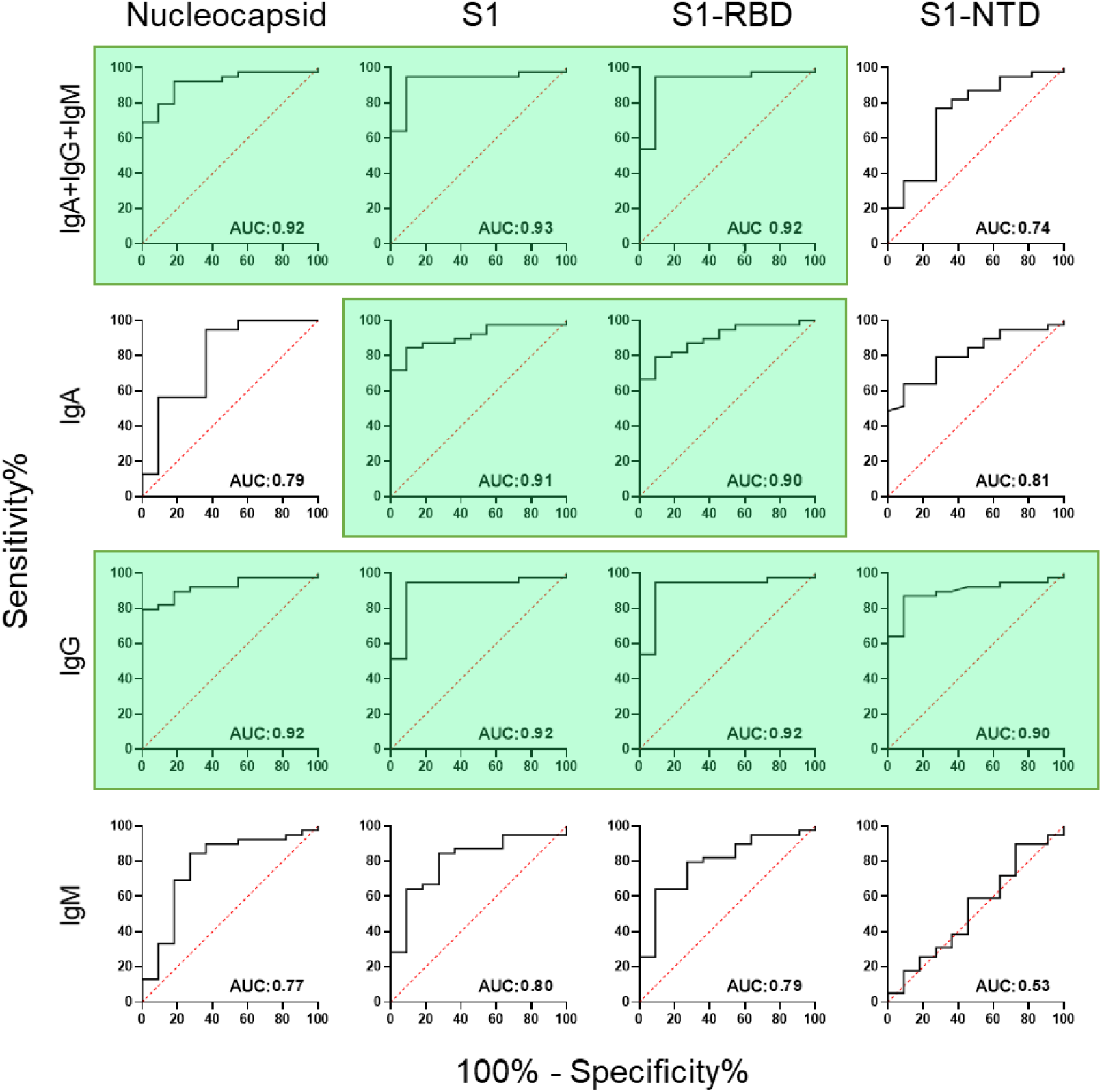
ROC curves distinguish severe/critical from mild/moderate SARS-CoV-2 infections during acute illness. SARS-CoV-2 antibody levels from the 39 severe/critical and 11 mild/moderate patients (Fig. 4) were examined using ROC curves. Samples were collected at least 6 DPSO and no more than 30 DPSO. AUC values near 0.5 indicate poor diagnostic potential (diagonal red line); AUC values near 1.0 reflect strong diagnostic value. Antigen:isotype combinations that yielded AUC values greater than 0.90 are highlighted in green.

**Table S1.** Demographics

| Subjects | **Convalescent**<br>**COVID-19+**<br>N= 69 | **Acute**<br>**COVID-19+**<br>N= 52 | **Pre-pandemic**<br>**Healthy Adults**<br>N= 106 | **Post-Pandemic**<br>**Healthy Adults**<br>N= 137 |
| --- | --- | --- | --- | --- |
| **Age, years**<br>(Mean ± SD) | 43 ± 15 | 52 ± 16 | 35 ± 12 | 33 ± 12 |
| **Sex, N (%)** |  |  |  |  |
| Males | 25 (36) | 31 (60) | 28 (26) | 63 (46) |
| Females | 44 (64) | 21 (40) | 78 (74) | 74 (54) |
| **Race, N (%)** |  |  |  |  |
| Caucasian | 35 (51) | 22 (42) | 35 (33) | 97 (71) |
| African American | 31 (45) | 26 (50) | 52 (49) | 9 (7) |
| Asian | 2 (3) | 4 (8) | 14 (13) | 20 (14) |
| Other | 1 (1) | 0 (0) | 5 (5) | 11 (8) |
| **Days post symptom onset**<br>Range Days<br>(Mean ± SD) | 67 ± 20 | 13 ± 6 | N/A | N/A |
| **COVID-19 severity**<br>**score<sup>28</sup>, N (%)** |  |  |  |  |
| Mild Illness | 34 (49) | 13 (25) |  |  |
| Moderate Illness | 16 (23) | 0 (0) | N/A | N/A |
| Severe Illness | 10 (15) | 3 (6) |  |  |
| Critical Illness | 9 (13) | 36 (69) |  |  |

**\*NIH management guidelines updated June 22, 2020**
**Asymptomatic** or pre-symptomatic infection: Individuals who test positive for SAR-CoV2 by viral PCR testing but have no symptoms.
**Mild Illness:** Individuals with any of the various signs and symptoms of COVID 19 (i.e. fever, cough, sore throat, malaise, headache, muscle pain) without shortness of breath, dyspnea, or abnormal chest imaging.
**Moderate Illness:** Individuals who have evidence of lower respiratory disease by clinical assessment of imaging and a saturation of oxygen (SpO<sub>2</sub>) ≥ 94% on room air at sea level.
**Severe Illness:** Individuals who have respiratory frequency > 30 breaths per minute, (SpO<sub>2</sub>) < 94% on room air at sea level, ratio of arterial partial pressure of oxygen to fraction of inspired oxygen (PaO<sub>2</sub>/FiO<sub>2</sub>) < 300 mmHg, or lung infiltrates > 50%.
**Critical Illness:** Individuals who have respiratory failure, septic shock, and/or multiple organ dysfunction.

**Table S2.** Antibody levels of severe/critical and mild/moderate SARS-CoV-2 groups converge overtime.

|  |  | 0-15 DPSO |  |  | 16-30 DPSO |  |  | 31-60 DPSO |  |  | 61-90 DPSO |  |  | 91-150 DPSO |  |  |
| --- | --- | --- | --- | --- | --- | --- | --- | --- | --- | --- | --- | --- | --- | --- | --- | --- |
|  |  | M/M<br>(N=13) | S/C<br>(N=33) | p-value | M/M<br>(N=6) | S/C<br>(N=23) | p-value | M/M<br>(N=21) | S/C<br>(N=9) | p-value | M/M<br>(N=21) | S/C<br>(N=9) | p-value | M/M<br>(N=25) | S/C<br>(N=5) | p-value |
| AGM | N | 8,642 | 58,972 | <.0001* | 8,132 | 53,924 | <.0001* | 13,375 | 36,588 | 0.0250* | 19,583 | 30,986 | 0.0785 | 12,757 | 12,696 | 0.9909 |
|  | S1 | 15,617 | 113,277 | <.0001* | 9,996 | 156,351 | <.0001* | 22,242 | 134,393 | 0.0032* | 36,163 | 112,541 | 0.0007* | 23,636 | 30,470 | 0.4385 |
|  | S1-RBD | 38,473 | 172,725 | <.0001* | 39,308 | 183,484 | <.0001* | 57,450 | 174,306 | 0.0001* | 77,756 | 194,211 | <.0001* | 66,512 | 88,412 | 0.2769 |
|  | S1-NTD | 2,154 | 3,963 | 0.0931 | 2,854 | 8,512 | 0.0201* | 1,432 | 23,591 | 0.1486 | 2,774 | 4,732 | 0.1169 | 1,729 | 2,794 | 0.2333 |
| IgA | N | 2,328 | 6,415 | 0.0523 | 182 | 4,595 | 0.0007* | 112 | 2,260 | 0.2794 | 101 | 583 | 0.1327 | 57 | 80 | 0.1791 |
|  | S1 | 3,182 | 15,126 | 0.0042* | 1,326 | 12,287 | <.0001* | 620 | 3,757 | 0.0500* | 1,842 | 3,186 | 0.4245 | 921 | 1,016 | 0.8964 |
|  | S1-RBD | 9,520 | 30,862 | 0.0102* | 9,099 | 21,472 | 0.0821 | 2,421 | 6,501 | 0.0697 | 6,463 | 12,406 | 0.4614 | 4,671 | 3,844 | 0.7868 |
|  | S1-NTD | 56 | 227 | 0.0005* | 69 | 637 | 0.0400* | 30 | 265 | 0.0815 | 10 | 55 | 0.0001* | 29 | 88 | 0.1786 |
| IgG | N | 2,827 | 36,091 | <.0001* | 3,781 | 35,539 | <.0001* | 9,783 | 24,622 | 0.0225* | 13,103 | 20,804 | 0.0989 | 7,419 | 8,396 | 0.8102 |
|  | S1 | 5,016 | 67,216 | <.0001* | 2,845 | 108,589 | <.0001* | 15,379 | 90,838 | 0.0038* | 21,499 | 77,841 | 0.0007* | 12,385 | 18,257 | 0.2395 |
|  | S1-RBD | 12,603 | 105,027 | <.0001* | 11,294 | 126,621 | <.0001* | 36,280 | 121,592 | 0.0001* | 46,986 | 141,427 | <.0001* | 37,308 | 57,409 | 0.1446 |
|  | S1-NTD | 77 | 1,593 | 0.0002* | 57 | 4,178 | <.0001* | 464 | 10,438 | 0.2496 | 618 | 1,997 | 0.0064* | 466 | 1,041 | 0.2077 |
| IgM | N | 1,849 | 5,842 | 0.0349* | 518 | 4,950 | 0.0062* | 662 | 2,380 | 0.3651 | 290 | 594 | 0.2224 | 278 | 235 | 0.6083 |
|  | S1 | 4,023 | 15,243 | 0.0063* | 2,128 | 20,111 | 0.0073* | 3,491 | 22,416 | 0.1373 | 1,705 | 7,751 | 0.0366* | 770 | 527 | 0.3154 |
|  | S1-RBD | 11,080 | 37,131 | 0.0026* | 6,921 | 42,051 | 0.0010* | 12,413 | 47,273 | 0.0886 | 6,570 | 22,125 | 0.0175* | 3,673 | 2,111 | 0.1264 |
|  | S1-NTD | 904 | 698 | 0.4444 | 1,160 | 1,470 | 0.5790 | 404 | 5,456 | 0.2753 | 686 | 734 | 0.8663 | 357 | 440 | 0.4527 |

Average net MFI was calculated for mild/moderate and severe/critical patients in 15-, 30-, and 60-day intervals over a five-month period post symptom onset. T-tests of significance were performed to compare the two groups at each time interval. Significant differences decreased between the two groups over time.

## References

1. J. F. Chan et al., Improved molecular diagnosis of COVID-19 by the novel, highly sensitive and specific COVID-19-RdRp/Hel real-time reverse transcription-polymerase chain reaction assay validated in vitro and with clinical specimens. Journal of clinical microbiology, (2020).

2. H. S. Wu et al., Serologic and molecular biologic methods for SARS-associated coronavirus infection, Taiwan. Emerg Infect Dis 10, 304–310 (2004).

3. J. Zhao et al., Antibody responses to SARS-CoV-2 in patients of novel coronavirus disease 2019. Clinical infectious diseases: an official publication of the Infectious Diseases Society of America, (2020).

4. R. C. Group et al., Dexamethasone in Hospitalized Patients with Covid-19 - Preliminary Report. The New England journal of medicine, (2020).

5. J. Grein et al., Compassionate Use of Remdesivir for Patients with Severe Covid-19. The New England journal of medicine 382, 2327–2336 (2020).

6. G. den Hartog et al., SARS-CoV-2-Specific Antibody Detection for Seroepidemiology: A Multiplex Analysis Approach Accounting for Accurate Seroprevalence. The Journal of infectious diseases 222, 1452–1461 (2020).

7. D. F. Gudbjartsson et al., Humoral Immune Response to SARS-CoV-2 in Iceland. The New England journal of medicine, (2020).

8. B. Isho et al., Persistence of serum and saliva antibody responses to SARS-CoV-2 spike antigens in COVID-19 patients. Sci Immunol 5, (2020).

9. N. Pisanic et al., COVID-19 serology at population scale: SARS-CoV-2-specific antibody responses in saliva. Journal of clinical microbiology, (2020).

10. Q. X. Long et al., Antibody responses to SARS-CoV-2 in patients with COVID-19. Nat Med 26, 845–848 (2020).

11. K. L. Lynch et al., Magnitude and kinetics of anti-SARS-CoV-2 antibody responses and their relationship to disease severity. Clinical infectious diseases: an official publication of the Infectious Diseases Society of America, (2020).

12. M. C. Woodruff et al., Extrafollicular B cell responses correlate with neutralizing antibodies and morbidity in COVID-19. Nature immunology, (2020).

13. M. S. Suthar et al., Rapid Generation of Neutralizing Antibody Responses in COVID-19 Patients. Cell Rep Med 1, 100040 (2020).

14. V. K. Shah, P. Firmal, A. Alam, D. Ganguly, S. Chattopadhyay, Overview of Immune Response During SARS-CoV-2 Infection: Lessons From the Past. Frontiers in immunology 11, 1949 (2020).

15. S. J. Zost et al., Rapid isolation and profiling of a diverse panel of human monoclonal antibodies targeting the SARS-CoV-2 spike protein. Nat Med 26, 1422–1427 (2020).

16. L. Liu et al., Potent neutralizing antibodies against multiple epitopes on SARS-CoV-2 spike. Nature 584, 450–456 (2020).

17. C. Kreer et al., Longitudinal Isolation of Potent Near-Germline SARS-CoV-2-Neutralizing Antibodies from COVID-19 Patients. Cell 182, 1663–1673 (2020).

18. E. Seydoux et al., Analysis of a SARS-CoV-2-Infected Individual Reveals Development of Potent Neutralizing Antibodies with Limited Somatic Mutation. Immunity 53, 98–105 e105 (2020).

19. W. Zhang et al., Molecular and serological investigation of 2019-nCoV infected patients: implication of multiple shedding routes. Emerg Microbes Infect 9, 386–389 (2020).

20. Z. Yongchen et al., Different longitudinal patterns of nucleic acid and serology testing results based on disease severity of COVID-19 patients. Emerg Microbes Infect 9, 833–836 (2020).

21. D. F. Robbiani et al., Convergent antibody responses to SARS-CoV-2 in convalescent individuals. Nature 584, 437–442 (2020).

22. L. Kuri-Cervantes et al., Comprehensive mapping of immune perturbations associated with severe COVID-19. Sci Immunol 5, (2020).

23. T. J. Ripperger et al., Detection, prevalence, and duration of humoral responses to SARS-CoV-2 under conditions of limited population exposure. medRxiv, (2020).

24. L. Zhang et al., Antibody responses against SARS coronavirus are correlated with disease outcome of infected individuals. J Med Virol 78, 1–8 (2006).

25. M. Dogan et al., Novel SARS-CoV-2 specific antibody and neutralization assays reveal wide range of humoral immune response during COVID-19. medRxiv, (2020).

26. A. Algaissi et al., SARS-CoV-2 S1 and N-based serological assays reveal rapid seroconversion and induction of specific antibody response in COVID-19 patients. Sci Rep 10, 16561 (2020).

27. A. S. Iyer et al., Dynamics and significance of the antibody response to SARS-CoV-2 infection. medRxiv, (2020).

28. C.-T. G. Panel. (National Institutes of Health, 2020).

29. C. W. Tan et al., A SARS-CoV-2 surrogate virus neutralization test based on antibody-mediated blockage of ACE2-spike protein-protein interaction. Nat Biotechnol 38, 1073–1078 (2020).

30. D. Mathew et al., Deep immune profiling of COVID-19 patients reveals distinct immunotypes with therapeutic implications. Science 369, (2020).

31. R. Varnaite et al., Expansion of SARS-CoV-2-Specific Antibody-Secreting Cells and Generation of Neutralizing Antibodies in Hospitalized COVID-19 Patients. Journal of immunology, (2020).

32. M. Heidelberger, F. E. Kendall, A Quantitative Study of the Precipitin Reaction between Type Iii Pneumococcus Polysaccharide and Purified Homologous Antibody. The Journal of experimental medicine 50, 809–823 (1929).

33. D. Bray, S. Lay, Computer-based analysis of the binding steps in protein complex formation. Proceedings of the National Academy of Sciences of the United States of America 94, 13493–13498 (1997).

34. H. Zinsser, Infection and Resistance. (The Macmillian Company, New York, NY, 1914).

35. G. Yan, D. Xing, S. Tan, Q. Chen, Rapid and sensitive immunomagnetic-electrochemiluminescent detection of p53 antibodies in human serum. Journal of immunological methods 288, 47–54 (2004).

36. G. D. Victora, M. C. Nussenzweig, Germinal centers. Annu Rev Immunol 30, 429–457 (2012).

37. J. A. Roco et al., Class-Switch Recombination Occurs Infrequently in Germinal Centers. Immunity 51, 337–350.e337 (2019).

38. N. Kaneko et al., Loss of Bcl-6-Expressing T Follicular Helper Cells and Germinal Centers in COVID-19. Cell 183, 143–157.e113 (2020).

39. M. Yuan et al., A highly conserved cryptic epitope in the receptor binding domains of SARS-CoV-2 and SARS-CoV. Science 368, 630–633 (2020).

